# Small-Molecule Inhibitors Allosterically Control Bruton’s Tyrosine Kinase Assembly on the Membrane

**DOI:** 10.64898/2026.10.08.757093

**Authors:** Alexander D. Fenton, Kathryn Todd, Shweta Shree, David Yin-wei Lin, Amy H. Andreotti, Jean K. Chung

## Abstract

Bruton’s tyrosine kinase (Btk) is targeted by small-molecule active-site inhibitors used to treat lymphomas and leukemias. Beyond its catalytic activity, Btk participates in signaling assemblies through scaffolding functions that can contribute to drug resistance. Here, we show that Btk forms micron-scale assemblies on membranes and that this behavior is allosterically regulated by ligand binding at the active site. Using supported lipid bilayers and total internal reflection fluorescence microscopy, we find that nucleotides and inhibitors including ibrutinib, acalabrutinib, and dasatinib promote assembly, whereas fenebrutinib and pirtobrutinib do not. Membrane assembly is independent of catalysis but correlates with ligand-dependent changes in Btk conformational flexibility measured by hydrogen-deuterium exchange. These results reveal a previously unrecognized connection between active-site ligand binding and mesoscale membrane organization, suggesting a mechanism by which kinase inhibitors may differentially affect Btk scaffolding functions implicated in drug resistance.

## Main Text

Bruton’s tyrosine kinase (Btk) is a central component in B-cell receptor signaling and an important drug target in B-cell malignancies.^1^ Multiple mechanistically distinct inhibitors are now in clinical use or in development, including covalent inhibitors,^2^ reversible inhibitors,^3^ and targeted degraders.^4^ While acquired resistance frequently arises through mutations in Btk that prevent efficient drug binding at the active site,^5^ more recently identified resistance mutations present a different problem: they impair Btk catalytic activity but still promote B-cell receptor signaling.^4^ These observations have drawn attention to Btk’s function as an organizer of signaling on the membrane,^6-11^ and how such functions are regulated independently of kinase activity. Whether active-site inhibitors themselves change these scaffolding functions is not known.

Btk contains an N-terminal pleckstrin homology-Tec homology (PHTH) module, followed by a proline-rich region (PRR), Src homology 3 and 2 (SH3, SH2) domains, and a C-terminal kinase domain (**Figure 1A**). The PHTH module recruits Btk to the plasma membrane through binding to phosphatidylinositol (3,4,5)-trisphosphate (PIP_3_).^12-15^ Once at the membrane, Btk can self-associate and activate via *trans*-autophosphorylation.^16,17^ We previously reported that truncated PHTH-PRR-SH3 constructs form membrane assemblies and that assembly requires two interfaces: the Saraste dimer formed by the PHTH module and the interaction between PRR and SH3 domain.^16^ The PHTH module and the kinase domain form at least transient direct interactions in solution, and that contact is relieved when Btk binds PIP_3_ at the membrane.^18-20^ Whether the kinase domain is also allosterically coupled to these assembly interfaces remains unknown. Such coupling would provide a mechanism by which ligand binding at the kinase active site could alter the mesoscale organization of Btk on the membrane. Consistent with this possibility, hydrogen-deuterium exchange-mass spectrometry (HDX-MS) studies of Btk have revealed ligand-dependent changes in conformational dynamics extending beyond the kinase domain.^5,21^ Long-range allosteric effects of ATP-competitive inhibitors are also increasingly recognized across the kinase family.^22^

**Figure 1.**
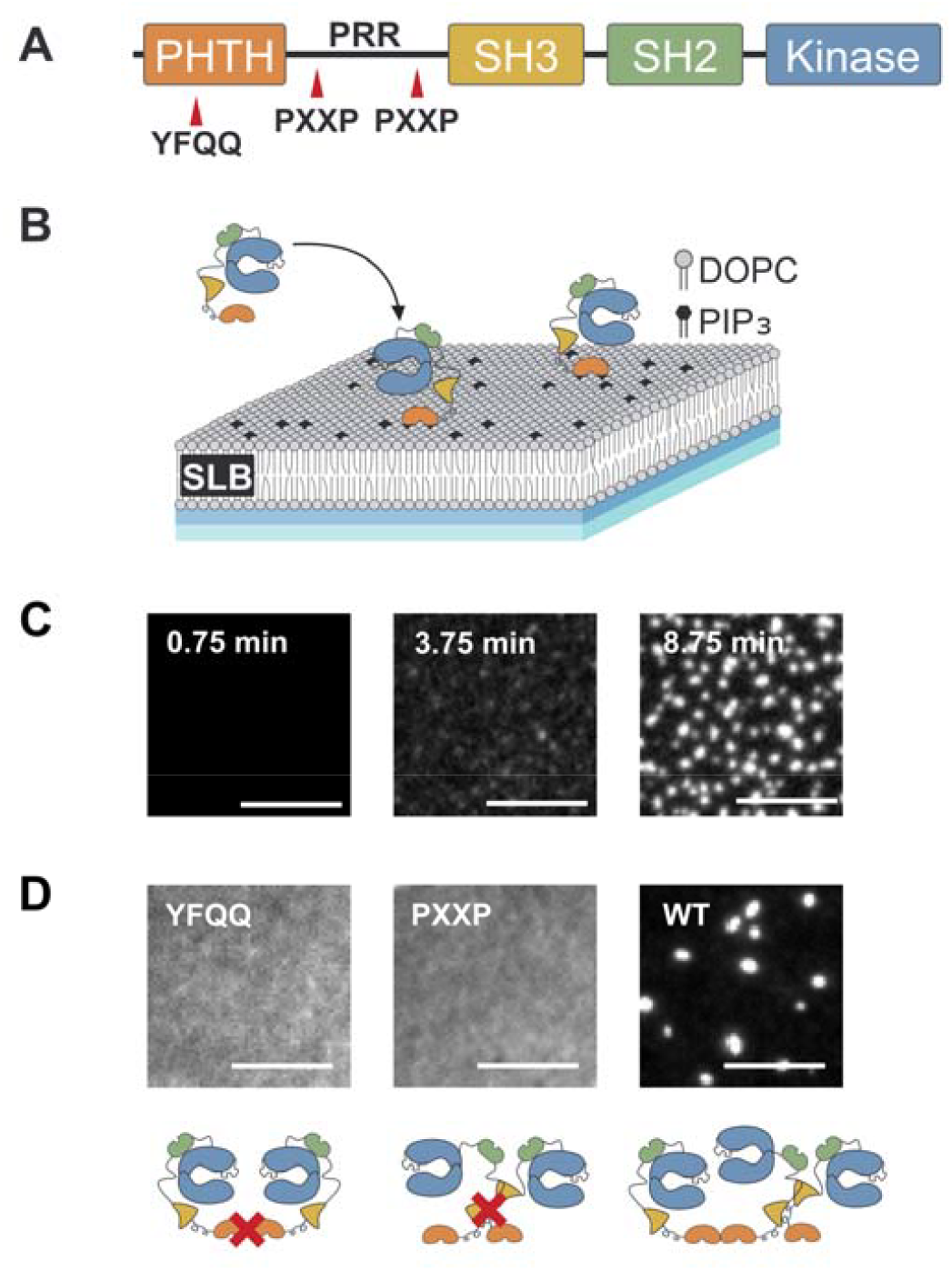
(**A**) Domain architecture of full-length Btk with the YFQQ and PXXP interface mutations indicated. (**B**) A schematic of the Btk reconstitution on supported lipid bilayers (SLBs). (**C**) Time series of Btk assembly with the same field of view shown at 0.75, 3.75, and 8.75 min after the addition of 200 nM WT Btk. (**D**) YFQQ, PXXP, and WT Btk more than 15 min after addition, at surface densities of 3,400, 3,400, and 1,200 Btk molecules/μm^2^ respectively. Diagrams below show the protein-protein interface disrupted by the indicated mutation. All experiments were performed on 4 mol % PIP_3_ SLBs with 1 mM ATP and 5 mM MgCl_2_, pH 7.4, at room temperature. Scale bars, 5 μm.

Here, we show that full-length Btk forms micron-scale assemblies on PIP_3_ membranes and that assembly is regulated by ligand binding at the kinase active site. Different active-site ligands produce distinct membrane organization, which correlates with changes in Btk conformational dynamics measured by HDX-MS. These results reveal a long-range coupling between the kinase domain and distant assembly interfaces.

To examine Btk assemblies on membranes, full-length Btk labeled with Alexa Fluor 488 was reconstituted on supported lipid bilayers (SLBs) containing 4 mol % PIP_3_ and 96 mol % 1,2-dioleoyl-sn-glycero-3-phosphocholine (DOPC). In all experiments, nucleotides or inhibitors were incubated with Btk in solution before introducing the mixture to the SLBs (**Figure 1B**). Membrane-associated Btk was visualized by total internal reflection fluorescence (TIRF) microscopy. Following addition of Btk with ATP, membrane-associated fluorescence increased over time, and assemblies emerged as Btk accumulated on the membrane (**Figure 1C**). Because assembly occurs on the two-dimensional membrane surface, surface density (Btk molecules/µm^2^) provides the relevant concentration variable for comparing assembly behavior across conditions. We therefore converted membrane fluorescence intensity to Btk surface density using a fluorescence calibration curve (**Figure S1**) and compare assembly behavior as a function of measured surface density.

We tested whether full-length Btk assembly depends on the same two intermolecular interfaces previously identified in the PHTH-PRR-SH3 construct. First, we perturbed the PHTH module. Disrupting the Saraste dimer interface with the previously established Y42Q/F44Q (YFQQ) mutations^13,23^ abolished assembly (**Figure 1D**, left). Similarly, perturbation of the PRR/SH3 interface with P189A/P192A/P203A/P206A (PXXP) mutations, which remove both PxxP motifs in the PRR,^24-30^ also abolished assembly (**Figure 1D**, middle). Both mutants remained dispersed at surface densities approximately threefold higher than those at which wild-type Btk assembled (**Figure 1D**, right). Both the PHTH dimerization and the PRR/SH3 interactions are required for full-length Btk assembly on PIP_3_ SLBs.

Having established that assembly requires specific regulatory domain interfaces, we next asked whether ligand binding status of the kinase domain affects Btk membrane organization. Btk assembly depends strongly on active-site nucleotide occupancy. Apo Btk recruited to the membrane but remained dispersed, whereas the nucleotides ATP and ADP each promoted assembly (**Figure 2**). Although ATP-induced assemblies showed robust Y551 phosphorylation (**Figure S2**), ADP also promoted assembly despite its inability to support phosphotransfer.^20,31^ These observations demonstrate that Btk assembly does not require kinase catalysis.

**Figure 2.**
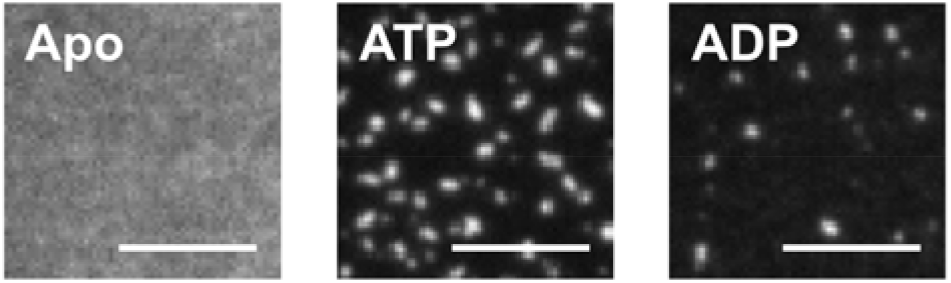
Active-site occupancy regulates Btk membrane organization. Btk on SLBs containing 4 mol % PIP_3_ in the absence of nucleotide (apo, 5,900 Btk molecules/µm^2^), in the presence of ATP (5,300 Btk molecules/µm^2^), or ADP (3,700 Btk molecules/µm^2^). Images were acquired 45 min after Btk addition. Nucleotides were at 1 mM and MgCl_2_ was at 5 mM. Scale bars, 5 µm.

Multiple clinically used Btk inhibitors act at the ATP-binding pocket. We therefore tested whether these inhibitors differentially affect membrane assembly. We compared two covalent inhibitors, ibrutinib and acalabrutinib, with three noncovalent inhibitors, dasatinib, fenebrutinib, and pirtobrutinib. All inhibitors were at 5 µM, approximately three orders of magnitude above published IC_50_ values for WT Btk.^32-35^ Anti-Btk(pY551) binding confirmed that Btk was catalytically active with ATP but inhibited by the inhibitors (**Figure S2**).

The inhibitor effects on Btk assembly were ligand-specific and did not correspond simply to covalent versus noncovalent binding. Dasatinib, ibrutinib, and acalabrutinib promoted Btk assembly, while Btk remained dispersed in the presence of fenebrutinib or pirtobrutinib (**Figure 3A**). Dasatinib produced extensive assemblies, whereas those formed with ibrutinib and acalabrutinib were comparatively sparse.

**Figure 3.**
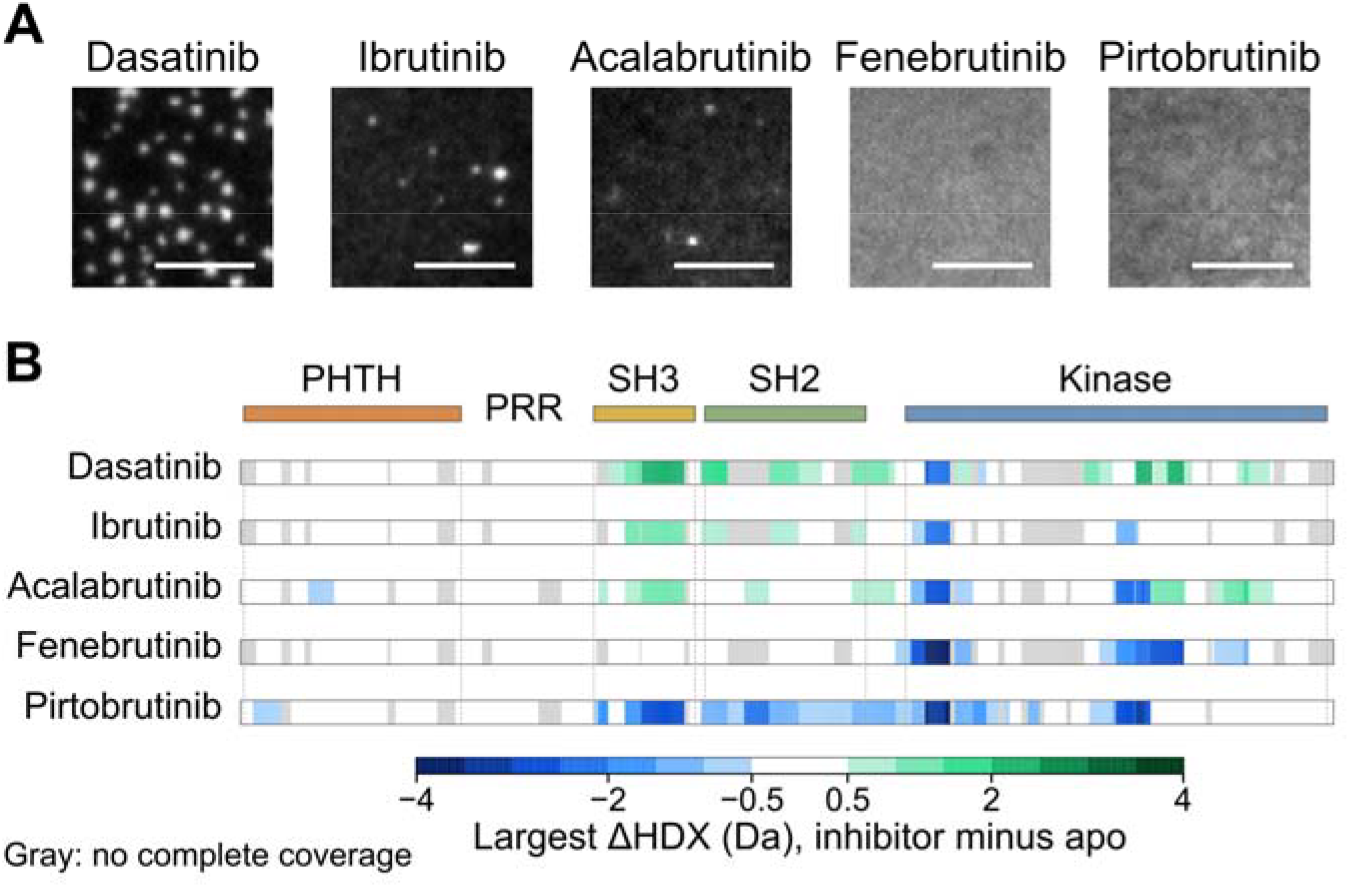
(**A**) Membrane organization of Btk in the presence of 5 µM dasatinib (6,200 Btk molecules/µm^2^), ibrutinib (7,300 Btk molecules/µm^2^), acalabrutinib (4,300 Btk molecules/µm^2^), fenebrutinib (10,000 Btk molecules/µm^2^), and pirtobrutinib (8,000 Btk molecules/µm^2^) with no added ATP on SLBs containing 4 mol % PIP_3_. Images taken 45 min after addition to the SLB. Scale bars, 5 µm. (**B**) Published inhibitor-minus-apo HDX differences from Joseph et al. 2020, 2024. A peptide was scored as changed if its difference reached 0.5 Da at one or more of five exchange times (10 s, 1 min, 10 min, 1 h, and 4 h), as defined by Joseph et al. Changed peptides are colored by their largest difference. At each sequence position, the color shows the median among overlapping peptides. Blue indicates less exchange and green more exchange. White indicates no change, and gray indicates no complete coverage. Drug order groups the responses for comparison and is not a quantitative ranking of global openness. Data are adapted from the cited eLife studies under CC BY; calculation details are in the SI.

The opposing effects of active-site inhibitors on Btk assembly suggested that different ligands stabilize distinct conformational states, consistent with the dynamic regulation of protein tyrosine kinases.^36^ Published HDX-MS measurements reveal ligand-dependent changes in amide protein exchange within the Btk SH3-SH2-kinase core (**Figures 3B** and **S3**).^5,21^ Binding of dasatinib, ibrutinib, and acalabrutinib causes an increase in deuterium uptake in portions of the SH3 and SH2 domains compared to apo Btk, while the kinase domain shows a mix of increases and decreases across these Btk-drug complexes. In contrast, binding of fenebrutinib and especially pirtobrutinib to full-length Btk results in protection from hydrogen/deuterium exchange. Inhibitors that increase exposure in the SH3-SH2 region are associated with Btk assembly, and inhibitors that either have no effect on deuterium exchange (fenebrutinib) or decrease the exchange in the SH3-SH2 region (pirtobrutinib) are associated with dispersed Btk. Assembly therefore tracks how each inhibitor changes Btk conformation and/or dynamics in solution and is not linked to the covalent versus non-covalent mechanism of drug action. HDX-MS data do not show significant effects of drug binding throughout the PHTH-PRR region of Btk likely due to the ‘fuzzy’^37^ conformational heterogeneity of the PHTH region^19^ and the preponderance of proline residues in the PRR resulting in fewer amide hydrogens.

To overcome the absence of signal for the PRR region in the HDX-MS experiment, we instead probed PRR accessibility directly on the membrane by monitoring recruitment of Grb2, an SH3-SH2-SH3 adaptor protein that binds the Btk PRR through its SH3 domains.^38^ Grb2 showed little binding to dispersed apo Btk but bound readily to ATP- and dasatinib-induced Btk assemblies (**Figure 4A**). This preferential recruitment is consistent with increased PRR accessibility in assembled Btk and supports a model in which assembly-promoting conditions favor PRR availability for intermolecular interactions (**Figure 4B**).

**Figure 4.**
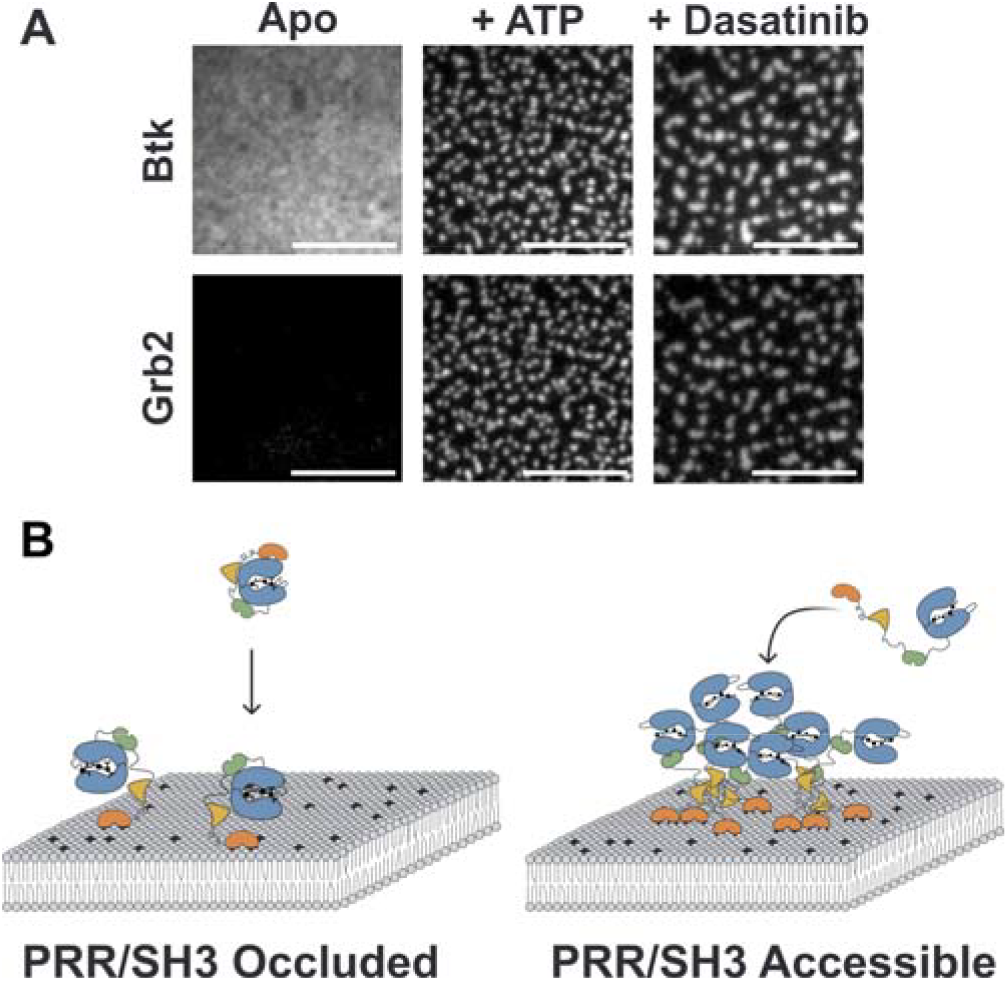
Assembly is associated with increased accessibility of the Btk PRR. (**A**) Btk-AF647 (top) and Grb2-AF555 (bottom) imaged in the same field on 4 mol % PIP_3_ SLBs under apo conditions or in the presence of ATP/MgCl_2_ (1 mM, 5 mM) or dasatinib (5 µM). Grb2 (20 nM) was added after Btk membrane recruitment an assembly. Scale bars, 5 µm. (**B**) Model for ligand-dependent regulation of Btk membrane assembly. Assembly-promoting conditions favor accessibility of the PRR-SH3 region, enabling intermolecular interactions.

Together, the results presented here show that active-site ligands regulate Btk membrane organization through long-range conformational coupling. HDX-MS measurements provide a structural basis for this effect. Remarkably, these solution-state conformational differences are reflected in Btk organization on the membrane, with increased hydrogen/deuterium exchange in the SH3-SH2 region of Btk correlating with increased assembly. An orthogonal membrane-based measurement further supports this model: Grb2 preferentially binds assembled Btk but not dispersed Btk, consistent with an available PRR in the assembled state. Thus, solution HDX-MS and membrane binding measurements converge on a model in which active-site ligands control the availability of distal interaction interfaces that drive Btk assembly.

Importantly, different inhibitors produce distinct membrane states despite targeting the same kinase active site. Because assembled Btk recruits Grb2 through its PRR more readily than dispersed Btk, the identity of inhibitor may determine whether Btk keeps its scaffolding interactions on the membrane even when catalytic activity is blocked. This distinction is relevant to drug resistance, since resistance mutations that impair Btk catalysis can still sustain B-cell receptor signaling.^4^ Whether inhibitors that prevent assembly also limit this scaffolding-dependent resistance remains to be determined. Together, our findings establish that active-site inhibitors regulate Btk not only through catalytic inhibition but also by controlling its membrane organization.^22^

## Associated Content

Supporting Information – Experimental methods, surface density calibration, pY551 antibody staining, and HDX structural mapping (PDF).

## Author Information

### Notes

The authors declare no competing financial interest.

## Acknowledgment

This work was supported by the National Science Foundation Grant No. 2238109.

## Table of Contents

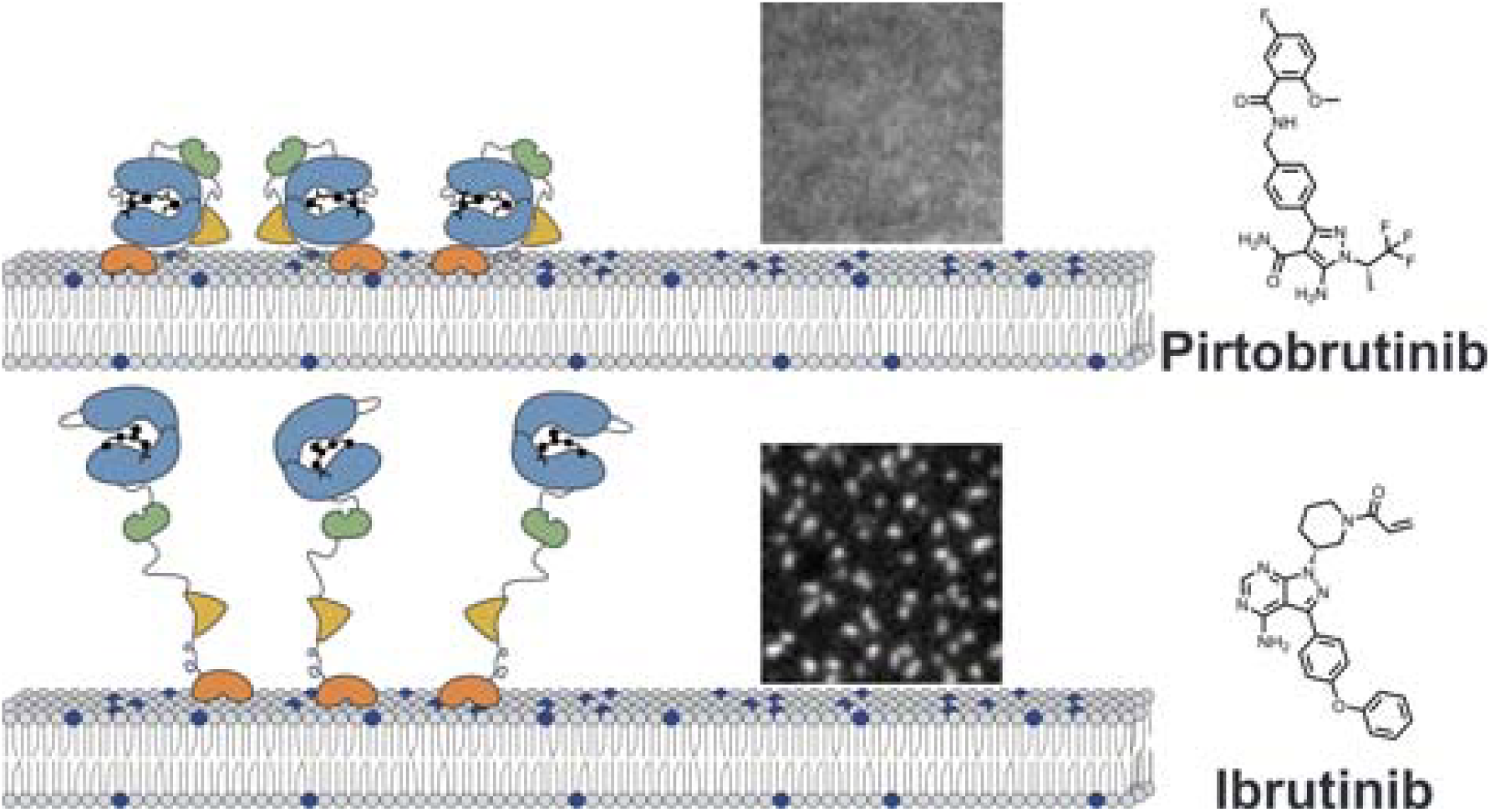

## Notes

### Competing Interest Statement

The authors have declared no competing interest.

## References

1 Hendriks, R. W., Yuvaraj, S. & Kil, L. P. Targeting Bruton’s tyrosine kinase in B cell malignancies. Nat Rev Cancer 14, 219–232 (2014). 10.1038/nrc3702

2 Honigberg, L. A. et al. The Bruton tyrosine kinase inhibitor PCI-32765 blocks B-cell activation and is efficacious in models of autoimmune disease and B-cell malignancy. Proc Natl Acad Sci U S A 107, 13075–13080 (2010). 10.1073/pnas.1004594107

3 Mato, A. R. et al. Pirtobrutinib in relapsed or refractory B-cell malignancies (BRUIN): a phase 1/2 study. Lancet 397, 892–901 (2021). 10.1016/S0140-6736(21)00224-5

4 Montoya, S. et al. Kinase-impaired BTK mutations are susceptible to clinical-stage BTK and IKZF1/3 degrader NX-2127. Science 383, eadi5798 (2024). 10.1126/science.adi5798

5 Joseph, R. E. et al. Impact of the clinically approved BTK inhibitors on the conformation of full-length BTK and analysis of the development of BTK resistance mutations in chronic lymphocytic leukemia. Elife 13 (2024). 10.7554/eLife.95488

6 Su, X. et al. Phase separation of signaling molecules promotes T cell receptor signal transduction. Science 352, 595–599 (2016). 10.1126/science.aad9964

7 Case, L. B., Zhang, X., Ditlev, J. A. & Rosen, M. K. Stoichiometry controls activity of phase-separated clusters of actin signaling proteins. Science 363, 1093–1097 (2019). 10.1126/science.aau6313

8 Huang, W. Y. et al. Phosphotyrosine-mediated LAT assembly on membranes drives kinetic bifurcation in recruitment dynamics of the Ras activator SOS. Proc Natl Acad Sci U S A 113, 8218–8223 (2016). 10.1073/pnas.1602602113

9 Sun, S., GrandPre, T., Limmer, D. T. & Groves, J. T. Kinetic frustration by limited bond availability controls the LAT protein condensation phase transition on membranes. Sci Adv 8, eabo5295 (2022). 10.1126/sciadv.abo5295

10 Huang, W. Y. C., Chiang, H. K. & Groves, J. T. Dynamic Scaling Analysis of Molecular Motion within the LAT:Grb2:SOS Protein Network on Membranes. Biophys J 113, 1807–1813 (2017). 10.1016/j.bpj.2017.08.024

11 Duarte, D. P. et al. Btk SH2-kinase interface is critical for allosteric kinase activation and its targeting inhibits B-cell neoplasms. Nat Commun 11, 2319 (2020). 10.1038/s41467-020-16128-5

12 Varnai, P., Rother, K. I. & Balla, T. Phosphatidylinositol 3-kinase-dependent membrane association of the Bruton’s tyrosine kinase pleckstrin homology domain visualized in single living cells. J Biol Chem 274, 10983–10989 (1999). 10.1074/jbc.274.16.10983

13 Hyvonen, M. & Saraste, M. Structure of the PH domain and Btk motif from Bruton’s tyrosine kinase: molecular explanations for X-linked agammaglobulinaemia. EMBO J 16, 3396–3404 (1997). 10.1093/emboj/16.12.3396

14 Fukuda, M., Kojima, T., Kabayama, H. & Mikoshiba, K. Mutation of the pleckstrin homology domain of Bruton’s tyrosine kinase in immunodeficiency impaired inositol 1,3,4,5-tetrakisphosphate binding capacity. J Biol Chem 271, 30303–30306 (1996). 10.1074/jbc.271.48.30303

15 Salim, K. et al. Distinct specificity in the recognition of phosphoinositides by the pleckstrin homology domains of dynamin and Bruton’s tyrosine kinase. EMBO J 15, 6241–6250 (1996). 10.1002/j.1460-2075.1996.tb01014.x

16 Fenton, A. D. & Chung, J. K. Membrane-dependent assembly of Bruton’s tyrosine kinase mediated by the Proline-rich region and SH3 domain. Protein Sci 34, e70213 (2025). 10.1002/pro.70213

17 Rawlings, D. J. et al. Activation of BTK by a phosphorylation mechanism initiated by SRC family kinases. Science 271, 822–825 (1996).

18 Amatya, N. et al. Lipid-targeting pleckstrin homology domain turns its autoinhibitory face toward the TEC kinases. Proc Natl Acad Sci U S A 116, 21539–21544 (2019). 10.1073/pnas.1907566116

19 Lin, D. Y. et al. Conformational heterogeneity of the BTK PHTH domain drives multiple regulatory states. Elife 12 (2024). 10.7554/eLife.89489

20 Wang, Q. et al. Autoinhibition of Bruton’s tyrosine kinase (Btk) and activation by soluble inositol hexakisphosphate. Elife 4 (2015). 10.7554/eLife.06074

21 Joseph, R. E. et al. Differential impact of BTK active site inhibitors on the conformational state of full-length BTK. Elife 9 (2020). 10.7554/eLife.60470

22 Vaisar, D. & Ahn, N. G. Latent allosteric control of protein interactions by ATP-competitive kinase inhibitors. Curr Opin Struct Biol 89, 102935 (2024). 10.1016/j.sbi.2024.102935

23 Chung, J. K. et al. Switch-like activation of Bruton’s tyrosine kinase by membrane-mediated dimerization. Proc Natl Acad Sci U S A 116, 10798–10803 (2019). 10.1073/pnas.1819309116

24 Rawlings, D. J. et al. Mutation of unique region of Bruton’s tyrosine kinase in immunodeficient XID mice. Science 261, 358–361 (1993). 10.1126/science.8332901

25 Park, H. et al. Regulation of Btk function by a major autophosphorylation site within the SH3 domain. Immunity 4, 515–525 (1996).

26 Hansson, H., Okoh, M. P., Smith, C. I., Vihinen, M. & Hard, T. Intermolecular interactions between the SH3 domain and the proline-rich TH region of Bruton’s tyrosine kinase. FEBS Lett 489, 67–70 (2001). 10.1016/s0014-5793(00)02438-8

27 Hansson, H., Smith, C. I. & Hard, T. Both proline-rich sequences in the TH region of Bruton’s tyrosine kinase stabilize intermolecular interactions with the SH3 domain. FEBS Lett 508, 11–15 (2001). 10.1016/s0014-5793(01)03018-6

28 Yu, H. et al. Structural basis for the binding of proline-rich peptides to SH3 domains. Cell 76, 933–945 (1994). 10.1016/0092-8674(94)90367-0

29 Lim, W. A., Richards, F. M. & Fox, R. O. Structural determinants of peptide-binding orientation and of sequence specificity in SH3 domains. Nature 372, 375–379 (1994). 10.1038/372375a0

30 Laederach, A. et al. Competing modes of self-association in the regulatory domains of Bruton’s tyrosine kinase: intramolecular contact versus asymmetric homodimerization. Protein Sci 11, 36–45 (2002). 10.1110/ps.26702

31 von Raussendorf, F., de Ruiter, A. & Leonard, T. A. A switch in nucleotide affinity governs activation of the Src and Tec family kinases. Sci Rep 7, 17405 (2017). 10.1038/s41598-017-17703-5

32 Hantschel, O. et al. The Btk tyrosine kinase is a major target of the Bcr-Abl inhibitor dasatinib. Proc Natl Acad Sci U S A 104, 13283–13288 (2007). 10.1073/pnas.0702654104

33 Patel, V. et al. Comparison of Acalabrutinib, A Selective Bruton Tyrosine Kinase Inhibitor, with Ibrutinib in Chronic Lymphocytic Leukemia Cells. Clin Cancer Res 23, 3734–3743 (2017). 10.1158/1078-0432.CCR-16-1446

34 Crawford, J. J. et al. Stereochemical Differences in Fluorocyclopropyl Amides Enable Tuning of Btk Inhibition and Off-Target Activity. ACS Med Chem Lett 11, 1588–1597 (2020). 10.1021/acsmedchemlett.0c00249

35 Gomez, E. B. et al. Preclinical characterization of pirtobrutinib, a highly selective, noncovalent (reversible) BTK inhibitor. Blood 142, 62–72 (2023). 10.1182/blood.2022018674

36 Amatya, N., Lin, D. Y. & Andreotti, A. H. Dynamic regulatory features of the protein tyrosine kinases. Biochem Soc Trans 47, 1101–1116 (2019). 10.1042/BST20180590

37 Arbesu, M., Iruela, G., Fuentes, H., Teixeira, J. M. C. & Pons, M. Intramolecular Fuzzy Interactions Involving Intrinsically Disordered Domains. Front Mol Biosci 5, 39 (2018). 10.3389/fmolb.2018.00039

38 Nocka, L. M., Eisen, T. J., Iavarone, A. T., Groves, J. T. & Kuriyan, J. Stimulation of the catalytic activity of the tyrosine kinase Btk by the adaptor protein Grb2. Elife 12 (2023). 10.7554/eLife.82676

