## Supporting Information for "Small-Molecule Inhibitors Allosterically Control Bruton’s Tyrosine Kinase Assembly on the Membrane"

### Btk expression and purification

Full-length Btk and the PXXP and YFQQ mutants were expressed as N-terminal 6×His–SUMO fusions in E. coli BL21(DE3) cells together with YopH phosphatase (Addgene Plasmid #79749). Starter cultures were grown overnight in LB medium with the appropriate antibiotics and used to inoculate larger cultures. Cells were grown at 37 °C to an OD600 of approximately 0.4–0.6, induced with 0.1 mM IPTG, and incubated at 18 °C for 16–20 h.

Harvested cells were resuspended in 50 mM Tris-HCl, pH 8.0, 500 mM NaCl, and 10% glycerol. Lysozyme (0.5 mg/mL), DNase I, and MgCl_2_ were added, and cells were lysed by sonication on ice. The lysate was clarified by centrifugation at 4 °C and incubated with Ni-NTA resin for approximately 2 h at 4 °C. The resin was transferred to a gravity-flow column and washed with lysis buffer. Bound protein was incubated overnight at 4 °C with Ulp1 protease and 2 mM TCEP to remove the His–SUMO tag on the column.

Cleaved Btk was collected in lysis buffer supplemented with 50 mM imidazole, concentrated, and purified by size-exclusion chromatography on a HiLoad 16/600 Superdex 200 column in 20 mM Tris-HCl, pH 8.0, 150 mM NaCl, 10% glycerol, and 1 mM DTT. Protein-containing fractions were checked by SDS-PAGE, pooled, concentrated, flash-frozen in liquid nitrogen, and stored at −80 °C. All constructs carry a Y617P mutation for expression. The PXXP construct contains P189A/P192A/P203A/P206A substitutions, and YFQQ contains Y42Q/F44Q substitutions; numbering refers to full-length Btk.

**WT Btk**

MGSSHHHHHHGSSMASMSDSEVNQEAKPEVKPEVKPETHINLKVSDGSSEIFFKIKKTTPLRRLMEAFAKRQGKEMDSLRFLYDGIRIQADQTPEDLDMEDNDIIEAHREQIGGMAAVILESIFLKRSQQKKKTSPLNFKKRLFLLTVHKLSYYEYDFERGRRGSKKGSIDVEKITCVETVIPEKNPPPERQIPRRGEESSEMEQISIIERFPYPFQVVYDEGPLYVFSPTEELRKRWIHQLKNVIRYNSDLVQKYHPCFWIDGQYLCCSQTAKNAMGCQILENRNGSLKPGSSHRKTKKPLPPTPEEDQILKKPLPPEPTAAPISTTELKKVVALYDYMPMNANDLQLRKGEEYFILEESNLPWWRARDKNGQEGYIPSNYITEAEDSIEMYEWYSKHMTRSQAEQLLKQEGKEGGFIVRDSSKAGKYTVSVFAKSTGEPQGVIRHYVVCSTPQSQYYLAEKHLFSTIPELINYHQHNSAGLISRLKYPVSKQNKNAPSTAGLGYGSWEIDPKDLTFLKELGTGQFGVVKYGKWRGQYDVAIKMIREGSMSEDEFIEEAKVMMNLSHEKLVQLYGVCTKQRPIFIITEYMANGCLLNYLREMRHRFQTQQLLEMCKDVCEAMEYLESKQFLHRDLAARNCLVNDQGVVKVSDFGLSRYVLDDEYTSSVGSKFPVRWSPPEVLMYSKFSSKSDIWAFGVLMWEIYSLGKMPYERFTNSETAEHIAQGLRLPRPHLASERVYTIMYSCWHEKADERPSFKILLSNILDVMDEES

**PXXP Btk**

MGSSHHHHHHGSSMASMSDSEVNQEAKPEVKPEVKPETHINLKVSDGSSEIFFKIKKTTPLRRLMEAFAKRQGKEMDSLRFLYDGIRIQADQTPEDLDMEDNDIIEAHREQIGGMAAVILESIFLKRSQQKKKTSPLNFKKRLFLLTVHKLSYYEYDFERGRRGSKKGSIDVEKITCVETVIPEKNPPPERQIPRRGEESSEMEQISIIERFPYPFQVVYDEGPLYVFSPTEELRKRWIHQLKNVIRYNSDLVQKYHPCFWIDGQYLCCSQTAKNAMGCQILENRNGSLKPGSSHRKTKKPLAPTAEEDQILKKPLAPEATAAPISTTELKKVVALYDYMPMNANDLQLRKGEEYFILEESNLPWWRARDKNGQEGYIPSNYITEAEDSIEMYEWYSKHMTRSQAEQLLKQEGKEGGFIVRDSSKAGKYTVSVFAKSTGEPQGVIRHYVVCSTPQSQYYLAEKHLFSTIPELINYHQHNSAGLISRLKYPVSKQNKNAPSTAGLGYGSWEIDPKDLTFLKELGTGQFGVVKYGKWRGQYDVAIKMIREGSMSEDEFIEEAKVMMNLSHEKLVQLYGVCTKQRPIFIITEYMANGCLLNYLREMRHRFQTQQLLEMCKDVCEAMEYLESKQFLHRDLAARNCLVNDQGVVKVSDFGLSRYVLDDEYTSSVGSKFPVRWSPPEVLMYSKFSSKSDIWAFGVLMWEIYSLGKMPYERFTNSETAEHIAQGLRLPRPHLASERVYTIMYSCWHEKADERPSFKILLSNILDVMDEES

**YFQQ Btk**

MGSSHHHHHHGSSMASMSDSEVNQEAKPEVKPEVKPETHINLKVSDGSSEIFFKIKKTTPLRRLMEAFAKRQGKEMDSLRFLYDGIRIQADQTPEDLDMEDNDIIEAHREQIGGMAAVILESIFLKRSQQKKKTSPLNFKKRLFLLTVHKLSYYEQDQERGRRGSKKGSIDVEKITCVETVIPEKNPPPERQIPRRGEESSEMEQISIIERFPYPFQVVYDEGPLYVFSPTEELRKRWIHQLKNVIRYNSDLVQKYHPCFWIDGQYLCCSQTAKNAMGCQILENRNGSLKPGSSHRKTKKPLPPTPEEDQILKKPLPPEPTAAPISTTELKKVVALYDYMPMNANDLQLRKGEEYFILEESNLPWWRARDKNGQEGYIPSNYITEAEDSIEMYEWYSKHMTRSQAEQLLKQEGKEGGFIVRDSSKAGKYTVSVFAKSTGEPQGVIRHYVVCSTPQSQYYLAEKHLFSTIPELINYHQHNSAGLISRLKYPVSKQNKNAPSTAGLGYGSWEIDPKDLTFLKELGTGQFGVVKYGKWRGQYDVAIKMIREGSMSEDEFIEEAKVMMNLSHEKLVQLYGVCTKQRPIFIITEYMANGCLLNYLREMRHRFQTQQLLEMCKDVCEAMEYLESKQFLHRDLAARNCLVNDQGVVKVSDFGLSRYVLDDEYTSSVGSKFPVRWSPPEVLMYSKFSSKSDIWAFGVLMWEIYSLGKMPYERFTNSETAEHIAQGLRLPRPHLASERVYTIMYSCWHEKADERPSFKILLSNILDVMDEES

### Grb2 expression and purification

Full-length Grb2 was expressed as an N-terminal 6×His–SUMO fusion in E. coli BL21 cells. Cultures were grown in LB containing 50 µg/mL kanamycin at 37 °C, induced at an OD600 of approximately 0.5–0.7 with 0.5 mM IPTG, and incubated overnight at 16 °C. Cells were lysed by sonication in 20 mM Tris-HCl, pH 8.5, 500 mM NaCl, 20 mM imidazole, and 5% glycerol. Clarified lysate was loaded onto Ni-NTA resin, washed in the same buffer, and eluted with an imidazole gradient to 500 mM.

The eluate was exchanged into 20 mM Tris-HCl, pH 8.0, 150 mM NaCl, 5% glycerol, and 1 mM DTT for overnight Ulp1 cleavage at 4 °C. A second Ni-NTA step removed His-tagged material. The flow-through containing cleaved Grb2 was concentrated and was further purified by size-exclusion chromatography on a HiLoad 16/600 column (Cytiva) equilibrated in 50 mM HEPES pH 7.4, 150 mM NaCl, and 5% glycerol.

**Grb2**

MGSSHHHHHHGSSMASMSDSEVNQEAKPEVKPEVKPETHINLKVSDGSSEIFFKIKKTTPLRRLMEAFAKRQGKEMDSLRFLYDGIRIQADQTPEDLDMEDNDIIEAHREQIGGMEAIAKYDFKATADDELSFKRGDILKVLNEECDQNWYKAELNGKDGFIPKNYIEMKPHPWFFGKIPRAKAEEMLSKQRHDGAFLIRESESAPGDFSLSVKFGNDVQHFKVLRDGAGKYFLWVVKFNSLNELVDYHRSTSVSRNQQIFLRDIEQMPQQPTYVQALFDFDPQEDGELGFRRGDFIHVMDNSDPNWWKGACHGQTGMFPRNYVTPVNRNV

### Fluorescent labeling

Btk and Grb2 exchanged into DTT-free buffer were labeled at cysteine residues with maleimide-functionalized Alexa Fluor dyes (Thermo Fisher Scientific). Proteins were reduced with a minimum 10-fold molar excess of TCEP for 30 min, and then dye was added at a 5-fold molar excess. Reactions were incubated overnight at 4 °C, protected from light, and quenched with 10 mM DTT. Free dye was removed on a desalting column or by repeated dilution and centrifugal concentration. Degree of labeling (DoL) was calculated from absorbance at 280 nm and at the dye's absorbance maximum. The A280 values were corrected for dye absorbance at 280 nm. Btk was labeled with AF488 for surface-density measurements and with AF647 for reporter assays. Grb2 was labeled with AF555. The AF488 DoL values used in density calculations were 0.29 for WT Btk, 1.26 for PXXP Btk, and 0.78 for YFQQ Btk. The Grb2-AF555 DoL was 0.49.

### Supported lipid bilayer preparation

Bilayers contained 96 mol % DOPC and 4 mol % PIP_3_. DOPC was supplied in chloroform, and PIP_3_ was dissolved in chloroform:methanol:water (8:8:3) before mixing. The solvent was removed under vacuum to form a lipid film. Films were hydrated with ultrapure water to 0.5 mg/mL total lipid and vortexed to disperse the lipids. Small unilamellar vesicles were prepared by tip sonication on ice at 33% amplitude, using 10 s on/10 s off pulses in a 90 s sonication program. The suspension was centrifuged at 18,000 × g for 20 min at room temperature, and the supernatant was used for bilayer formation.

Glass coverslips were bath-sonicated in isopropanol and then water for 15 min each, etched with freshly prepared piranha solution (3:1 sulfuric acid:hydrogen peroxide), and rinsed thoroughly with ultrapure water. Coverslips were dried with nitrogen and mounted in ibidi six-channel sticky slides. Vesicles were diluted to 0.125 mg/mL in 0.75× PBS, and 100 µL was added per chamber. After 15 min at room temperature, each chamber was washed with 1 mL PBS, incubated with 1 mg/mL κ-casein for 10 min, and washed with another 1 mL PBS. Imaging buffer was PBS with 10 mM β-mercaptoethanol and 0.1 mg/mL κ-casein. Bilayers were used on the day of preparation.

### Membrane assembly assays

Btk, nucleotide, and inhibitor solutions were prepared in imaging buffer and mixed before loading onto bilayers. Samples were introduced through one chamber well while removing an equal volume from the opposite well. Experiments were performed at room temperature. ATP or ADP was used at 1 mM with 5 mM MgCl_2_. Inhibitors were diluted from DMSO stocks to 5 µM with 1% final DMSO. The five-inhibitor comparison in Figure 3 had no added ATP.

Figure 1C used 200 nM WT Btk. Figure 1D used 25 nM WT and 100 nM PXXP or YFQQ. The ADP condition in Figure 2 used 275 nM WT Btk. Figure 3 used 382 nM WT Btk for fenebrutinib and 100 nM for the other inhibitors. The higher concentration with fenebrutinib was used to reach a Btk surface density comparable to the other inhibitor conditions.

### Grb2 recruitment and antibody staining

For Grb2 recruitment, 100 nM WT Btk-AF647 was incubated on bilayers for 8 min. Grb2-AF555 was then introduced at 20 nM. ATP/MgCl_2_ or dasatinib was maintained in the corresponding reporter solution. Images were acquired approximately 4 min after Grb2 addition.

For antibody staining, membrane-bound WT Btk-AF647 was exposed to PE-conjugated anti-Btk pY551 antibody (Biolegend PE anti-BTK Phospho (Tyr551)/ITK Phospho (Tyr512)). Figure S2A used 100 nM Btk and a 2 µg/mL antibody flow solution under an ATP-containing condition. Figure S2B used 100 nM Btk and 1 µg/mL antibody with dasatinib in the presence of ATP. Paired Btk and antibody images were recorded every 15 s. The displayed frames were acquired 2.8 min (S2A) and 4.0 min (S2B) after recording began.

### TIRF microscopy

Images were acquired on a Nikon Eclipse Ti2 microscope with a 100× NA 1.45 oil-immersion objective and an Andor DU-897 EMCCD camera. AF488, AF555/PE, and AF647 were excited at 488, 561, and 640 nm, respectively. The pixel size was 0.16 µm.

### Surface-density calibration and image analysis

Btk surface density was estimated using Oregon Green 488-DHPE bilayers of known dye density (Figure S1). Five standards spanning 347.5–2,780 OG molecules/µm² were each imaged in three fields. Camera and blank-bilayer backgrounds were removed, and intensities were normalized for exposure and gain. A straight line fitted through zero gave a slope of 70 ADU per OG molecule/µm² at 25 ms exposure and EM gain 100. The measured brightness of AF488-Btk relative to OG was 0.095.

Mean intensity was measured from the raw pixels in each displayed crop. After background subtraction and exposure normalization, intensity was divided by the calibration slope, the relative-brightness factor, and the dye-to-protein ratio. A pre-protein frame from the same movie supplied the background when available; otherwise, a blank-bilayer image at the corresponding exposure was used. Densities are approximate estimates reported to two significant figures. The calibration was applied only to AF488-Btk, and values above the standard range require linear extrapolation.

Image brightness was adjusted linearly for display, after density measurements were made from the raw images. Paired channels use the same crop. All microscopy panels span 10.08 × 10.08 µm and have 5 µm scale bars.

### Hydrogen deuterium exchange analysis

Published inhibitor-minus-apo peptide measurements were obtained from Joseph et al. (2020), Figure 5 Source Data 1, and Joseph et al. (2024), Figure 4 Supplementary File 2 [1,2]. Only peptides measured at all five times were included. A peptide was scored as changed if its difference reached 0.5 Da at one or more times, the no-change criterion used by Joseph et al. Changed peptides take the signed difference from the time point with the largest change, and unchanged peptides were set to zero. Where peptides overlap, the median sets the color. Blue indicates less exchange, green more exchange, white no change, and gray no complete coverage. The strips summarize peptide-level data rather than measurements at individual residues. Domain spans were PHTH 1–170, PRR 171–213, SH3 214–274, SH2 281–377, and kinase 402–655. The PHTH and PRR profiles do not provide clear evidence of ligand-dependent PRR dynamics, as only a single PHTH peptide exceeded 0.5 Da with acalabrutinib or pirtobrutinib, and no PRR peptide did. The Grb2 recruitment measurements provide the experimental support for the PRR-accessibility model.

For Figure S3, the same values were displayed on a common SH3–SH2–kinase structural template in UCSF ChimeraX. A compact core was assembled from the biological assembly of PDB 4XI2. Only residues with coordinates were displayed. PHTH and PRR were omitted because they are absent from this structure. Color shows the HDX difference and tube width shows its magnitude; neither represents a measured change in the protein’s three-dimensional shape.

### References

[1] Joseph, R. E.; et al. Differential impact of BTK active site inhibitors on the conformational state of full-length BTK. eLife 2020, 9, e60470. https://doi.org/10.7554/eLife.60470

[2] Joseph, R. E.; et al. Impact of the clinically approved BTK inhibitors on the conformation of full-length BTK and analysis of the development of BTK resistance mutations in chronic lymphocytic leukemia. eLife 2024, 13, RP95488. https://doi.org/10.7554/eLife.95488.3

HDX data were adapted under the CC BY licenses of these articles.

### Supporting figures

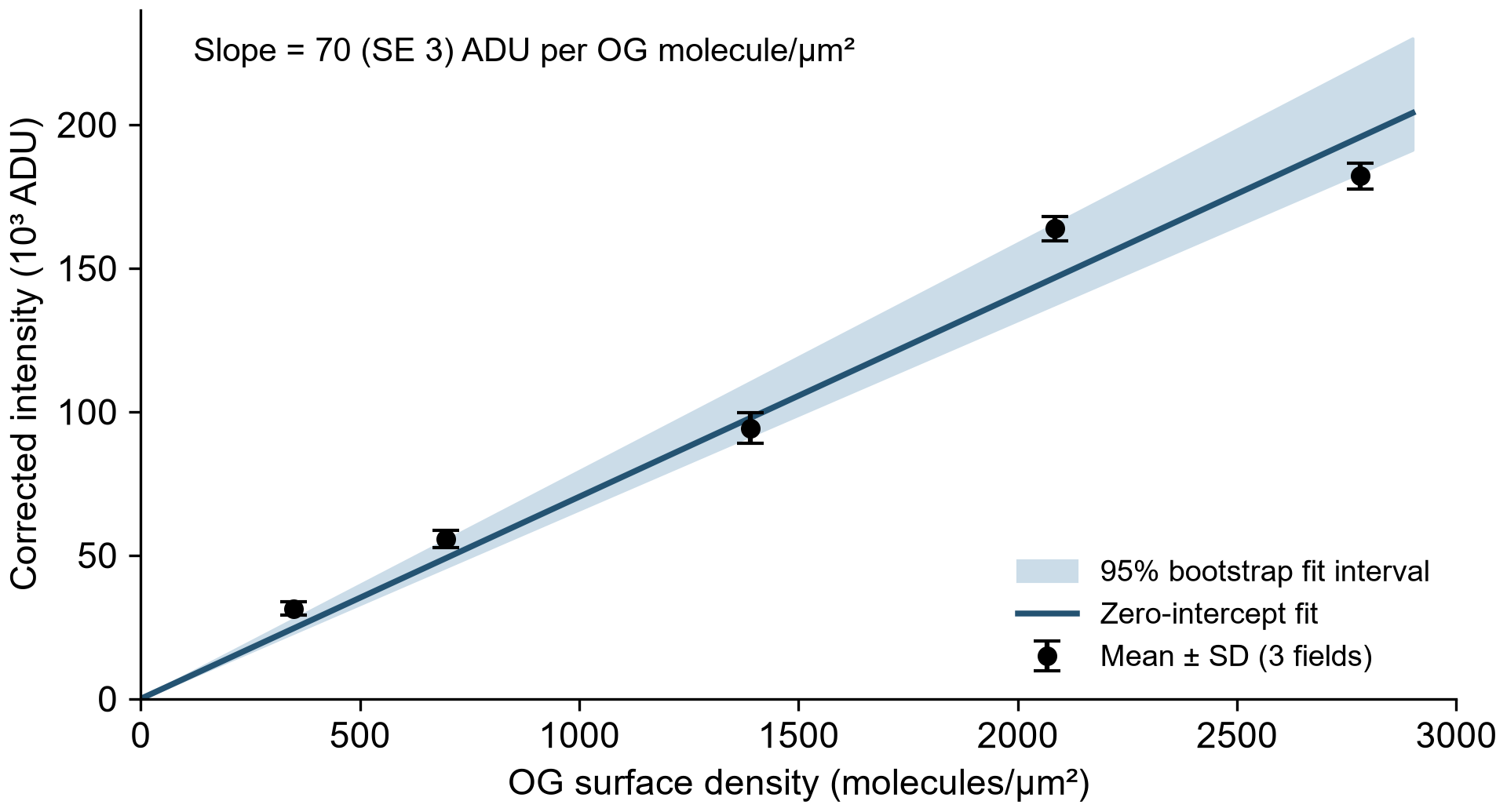

Figure S1. Surface-density calibration. Points show mean ± SD from three fields for each of five OG-DHPE standards. The line is a fit through zero with slope 70 ADU per OG molecule/µm². The shaded region shows the 95% bootstrap interval for the fitted line (slope, 66–79), calculated by resampling the five standard means. Intensities are background corrected and normalized to 25 ms exposure and EM gain 100.

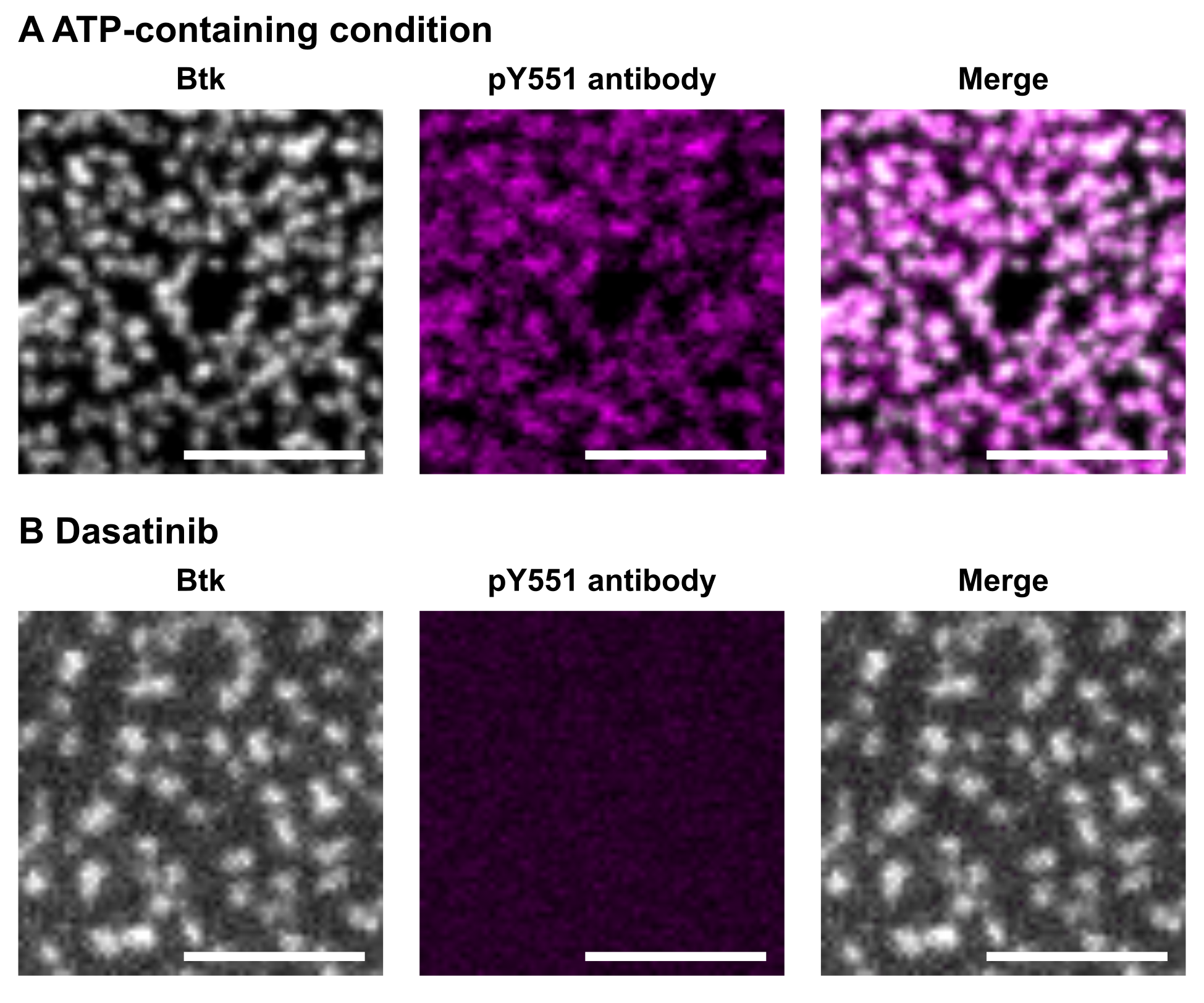

Figure S2. Y551 antibody staining of membrane-bound Btk. (A) An ATP-containing sample. (B) Dasatinib-treated Btk in the presence of ATP. Both samples contained 100 nM WT Btk-AF647 on 4 mol % PIP_3_ bilayers. Btk is shown in grayscale and PE-conjugated anti-Btk pY551 antibody in magenta. Images were acquired 2.8 min (A) or 4 min (B) after recording began. Scale bars, 5 µm.

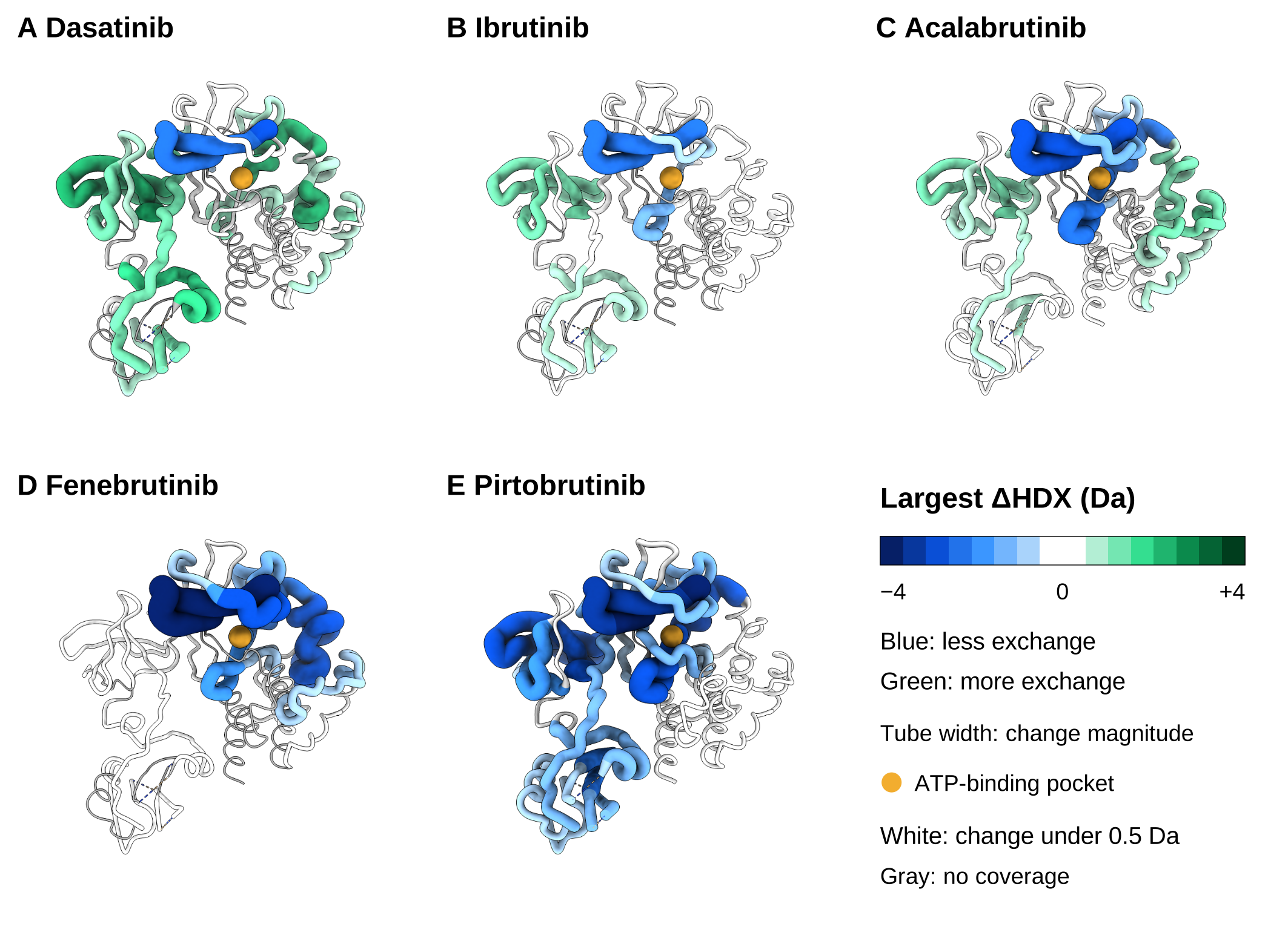

Figure S3. Structural context for inhibitor-dependent HDX changes. The values used in Figure 3B are mapped onto the Btk SH3–SH2–kinase core using PDB 4XI2. (A) Dasatinib. (B) Ibrutinib. (C) Acalabrutinib. (D) Fenebrutinib. (E) Pirtobrutinib. Blue indicates less exchange and green more exchange relative to apo Btk; tube width increases with the magnitude of the change. White indicates no change (less than 0.5 Da), and gray indicates missing HDX coverage. The gold sphere marks the approximate ATP-binding pocket, not a docked inhibitor. Each panel uses the same structural template; the shapes are not drug-specific conformations. PHTH and PRR are absent from these coordinates. Colors summarize overlapping peptide measurements and do not resolve individual residues. Data from Joseph et al. [1,2].
